# Stressostat cultivation of *Lactococcus lactis* improves lactate stress resistance through mutations in RNA polymerase

**DOI:** 10.64898/2026.04.14.718377

**Authors:** Sylviani Hartono, Henriette Lyng Røder, Sjef Boeren, Daan C. Swarts, Tjakko Abee, Eddy J. Smid, Oscar van Mastrigt

**Affiliations:** Food Microbiology, Wageningen University & Research, Wageningne, PO Box 17, 6700AA, The Netherlands; Section for Food Microbiology, Gut Health and Fermentation, Department of Food Science, University of Copenhagen, Frederiksberg 1958, Denmark; Laboratory of Biochemistry, Wageningen University & Research, Wageningen 6708WE, The Netherlands

**Keywords:** Lactic acid bacteria, starter culture, RNA polymerase, evolution, osmotic stress, arginine metabolism, *Lactococcus lactis*

## Abstract

Adaptive laboratory evolution is used to improve the phenotypes of microorganisms and to characterise the mechanisms underlying resistance against complex growth inhibition. Here we focused on lactic acid bacteria (LAB) as starter cultures for food fermentations. Production of LAB starter cultures is challenging due to growth inhibition by organic acids, mainly lactate, produced during fermentation. By utilising stressostat cultivation we generated *Lactococcus lactis* isolates with enhanced lactate resistance. Using a combination of (meta)genomics, proteomics and pH-controlled batch fermentations, we deciphered the lactate resistance mechanisms of these *L. lactis* isolates. Proteome responses of *L. lactis*, combined with similar growth inhibition at high salt, suggest that high lactate mainly causes osmotic stress. We identified RNA polymerase (RNAP) mutations in subunits β (*rpoB*) and β’ (*rpoC*) as key mutations, causing pleiotropic effects in the proteome. These proteome adaptations are linked to enhanced lactate resistance, particularly the resistance to hyperosmotic stress without glycine-betaine supplementation, likely by altering cross-linking of the peptidoglycan in the cell envelope via downregulation of MurE. The proteome changes indicate that lactate might (indirectly) cause oxidative stress. Combined, our study shows that RNAP mutations enhanced lactate resistance through pleotropic effects in the proteome that changed *L. lactis* responses against multiple stresses.

**Highlights:**

- Lactate resistant variants were isolated from stressostat cultivations
- Metagenomics revealed evolutionary trajectory during stressostat cultivation
- RNAP mutations were identified as key drivers for improved lactate resistance
- RNAP mutations altered the proteome, enhancing the osmotic stress resistance

## Introduction

Microorganisms experience rapid evolution to adapt to environmental changes due to their short generation time, large population size, and small genome size. This makes microorganisms ideal subjects for adaptive laboratory evolution (ALE) (Bennett and Hughes, 2009). ALE has been widely used to improve microbial phenotypes for industrial application and to investigate the emergence of complex traits (Bachmann et al., 2015; Mozhayskiy and Tagkopoulos, 2013; Sandberg et al., 2019). Combined with new generation sequencing and - omics technologies, ALE can be used as a top-down approach to investigate the molecular adaptation and underlying mechanism of microbial resistance against complex stresses (Bachmann et al., 2015; Brockhurst et al., 2011; Mozhayskiy and Tagkopoulos, 2013).

Lactic acid bacteria (LAB) populations are found in many fermented food products from dairy or plant sources. LAB are widely used as starter cultures to ensure the quality and safety of these fermented food products (Leroy and De Vuyst, 2004; Ghoddusi, 2011; Lee et al., 2020). The starter culture industry produces LAB in pH-controlled batch fermentations. Despite a surplus of substrates and control of the pH at optimal levels, growth of the LAB typically stops before substrates are depleted. It is hypothesized that their end-product, lactic acid, is the major growth limiting factor for LAB biomass production (Boonmee, 2010; Straathof, 2023), but it is unclear how lactate inhibits growth. At neutral pH, growth inhibition by passive diffusion of undissociated lactic acid and subsequent acidification of the cytoplasm is minimal as most of weak acids will be in the dissociated form (Gonçalves et al., 1997; Rault et al., 2009; Robergs et al., 2023; Carpenter and Broadbent, 2009). Instead, accumulation of dissociated lactic acid (lactate) and counter-ions required for pH control might increase the osmolarity of the growth medium and thereby could inhibit growth. In line with that hypothesis, a study by Loubierre et. al (Loubiere et al., 1997), showed that lactic acid and sodium chloride inhibited *Lactococcus lactis* to a similar extent, indicating that lactate accumulation indeed exerts osmotic stress. However, they observed discrepancy in the growth rate observed in batch culture with pH control compared to that observed without pH control, indicating that lactate could induce more than just osmotic stress in pH-controlled cultures. For instance, at high extracellular lactate concentrations, lactate might accumulate intracellularly, which could inhibit important metabolic pathways, disrupt energy generation, and/or lead to accumulation of toxic intermediates (Roe et al., 2002).

We previously generated *L. lactis* FM03P isolates with improved lactate resistance and biomass production in a pH-controlled batch fermentation using stressostat cultivation (Hartono et al., 2023). Stressostat cultivation is an ALE method that uses continuous cultivation in substate surplus to *in situ* increase the end-product concentration, resulting in a gradually increasing evolutionary pressure. In this way, stressostat selects for isolates with improved end-product resistance during cultivation. At high lactate concentrations, stressostat isolates grew faster than the wild-type (WT) isolates, indicating an improved lactate resistance (Hartono et al., 2023). Furthermore, some isolates, notably FM105, show prolonged biomass production at high lactate concentrations and improved survival in the stationary phase when grown in a pH-controlled batch fermentation (Hartono et al., 2023). However, it is not known how these isolates improved their resistance towards lactate.

In this study, we aimed at deciphering the lactate resistance mechanism of *L. lactis* stressostat isolates using a combination of pH-controlled batch fermentations, (meta)genomics, and proteomics. Metagenomics and whole-genome sequencing of the stressostat isolates revealed mutations in RNA polymerase that improved the lactate resistance of *L. lactis*. This results in divergent proteome profiles compared to wild-type *L. lactis* during pH-controlled batch cultivation. Finally, we show how these mutations and divergent proteomes change the phenotypes of the stressostat isolates.

## Materials & Methods

### Strain and pre-culture condition

*Lactococcus lactis* biovar diacetylactis FM03P (NCBI ASM214821v1) and its stressostat isolates (Hartono et al., 2023; van Mastrigt et al., 2017) were used in this study. For each cultivation, *L. lactis* was streaked on M17 agar plates (BD Difco, France) supplemented with 0.5% (w/v) lactose (LM17) and incubated at 30°C for 72 hours. A single colony was inoculated into LM17 broth and incubated overnight at 30°C. These overnight cultures were used for further experiments.

### DNA isolation

*L. lactis* FM03P cultures (evolved communities or overnight cultures of wild type or evolved isolates) were subjected to genome sequencing. For isolation of genomic DNA, cells were harvested from 1 ml culture by centrifugation at 17000×*g* for 2 min and washed 2 times with 1 ml peptone physiological salt (PPS, Tritium Microbiologie, The Netherlands). The cell pellets were resuspended in 180 µL lysis buffer consisting of 20 mM Tris-HCl pH 8, 2 mM EDTA, 1.2% (w/v) Triton X-100, and 20 mg ml^-1^ lysozyme, and incubated for 1 hour at 37°C. The extraction procedure was continued by following the protocol of the DNeasy Blood and Tissue Kit (Qiagen, Germany). The extracted DNA was further concentrated/purified by adding 0.1× volume of 3 M Na-Acetate pH 5.2, 1× volume of isopropanol and incubated at -20°C for 20 mins. The DNA pellet was extracted by centrifugation (17000×g for 15 min at 4°C) and washed with 70% ethanol. The ethanol was removed by centrifugation (17000× g for 15 min at 4°C) and air drying for 15 min. The DNA pellet was resuspended in 100 µL milliQ water.

### Metagenomic analysis

The isolated DNA of wild type *L. lactis* FM03P and evolved communities were sequenced by using Illumina NovaSeq 6000 system (Eurofins Genomics, Germany) with 2 × 150 bp paired-end read mode and a total of 10 million read pairs. The metagenome sequencing results are deposited under study PRJEB108644 in the European Nucleotide Archive (ENA) of EMBL-EBI. The RAW reads were analysed using breseq (v0.35.0) (Deatherage and Barrick, 2014) to call variants with the—p flag, depending on the case. This analysis was performed against the reference genome, *Lactococcus lactis* bv. diacetylactis strain FM03 (NCBI ASM214821v1) and mutations found in our ancestral strain compared to the reference genome were not taken into account. Additionally, we filtered and showed only the mutations that appeared in at least 5% of the reads for the metagenomic analysis.

### Whole genome sequencing and SNP analysis

The DNA was sequenced by using Illumina NovaSeq 6000 system (Eurofins Genomics, Germany) with 2 × 150 bp paired-end read mode and minimum 100× average coverage. The whole genome sequencing results are deposited under study PRJEB108642 in the European Nucleotide Archive (ENA) of EMBL-EBI. Raw reads were processed using Galaxy platform (The Galaxy Community, 2022) and trimmed with cutadapt (Martin, 2011). Snippy (Seemann, 2015) was used for SNP analysis of the isolates against FM03P as reference, with a minimum proportion for variant evidence at 95%.

### Proteomic analysis

Proteomic analysis was performed on pH-controlled batch cultures of FM03P, FM105, and FM211. Cultures were grown in chemically defined media as described before (Hartono et al., 2023) with omission of the nucleotides stock solution. The conditions were set as follows: temperature at 30°C, stirring speed at 200 rpm, and pH at 6.5. The pH was maintained by automatic addition of 10 M KOH. In addition, the headspace was flushed with nitrogen gas at a flow of 0.1 L min-1 to maintain anaerobic condition. Samples were taken at the exponential phase (lactate ∼100 mM) and at the beginning of the stationary phase (lactate 600-700 mM, depending on the isolates). From each sampling point, 2 ml culture was pelleted by centrifugation (17000×g for 2 mins at 4°C) and stored at -80°C. The pellets were thawed on ice, washed 2 times with 100 mM Tris (pH 8), and resuspended in Tris buffer with Pierce^TM^ protease inhibitor cocktail (ThermoFisher Scientific, USA) and 15 mM dithiothreitol. Cells were lysed by sonication twice for 45 s with 30 s rest on ice in between.

The protein aggregation capture (PAC) method (Batth et al., 2019) was used in a modified way for the preparation of samples for proteomic analysis. In 100 µl sonicated sample, proteins were reduced by adding 0.1 volume 150 mM dithiothreitol (DTT) and incubating at 45°C for 30 mins. Subsequently, proteins were unfolded by adding 3 volumes freshly prepared 8 M urea in 100 mM Tris (pH8), and alkylated by adding 0.1 volume 200 mM acrylamide followed by incubation at room temperature for 30 mins. The pH was then adjusted to pH 7 by adding 7 µl 10% trifluoro-acetic acid (TFA). Two different 50 µg µl^-1^ SpeedBeads (magnetic carboxylate modified particles, GE Healthcare, USA) of product number 45152105050250 + 65152105050250 were mixed in 1:1 ratio and 8 µl was added into each sample. 2.5 volumes acetonitrile was added into the sample-beads mixture. The samples were incubated with gentle shaking at room temperature for 20 mins. The beads containing protein were separated from the supernatant using a magnetic rack. The beads were then washed sequentially with 1 ml 70% ethanol and 1 ml 100% acetonitrile and resuspended in 100 µl trypsin solution (5 ng µl^-1^ in 50 mM ammonium bicarbonate) and incubated overnight with gentle shaking at room temperature. The pH was then adjusted to pH 3 by adding 4 µl 10% TCA to stop trypsin activity. The beads were separated from the peptides solution using magnetic racks and washed with 100 µl formic acid (1 ml l^-1^) to elute the remaining peptides from the beads. All eluents were combined and concentrated until 10-15 µl volume. The end volume was adjusted to 50 µl with 1 ml l^-1^ formic acid. A maximum of 5 µL sample was injected into nLC-MSMS as described before (Liu et al., 2021).

LC-MS data analysis (false discovery rates were set to 0.01 on peptide and protein levels) and additional result filtering (minimally 2 peptides are necessary for protein identification of which at least one is unique and at least one is unmodified) were performed as described previously(Zheng et al., 2024). *Lactococcus lactis* bv. diacetylactis strain FM03 (NCBI ASM214821v1) database was used as protein sequences reference. The nLC-MSMS system quality was checked with PTXQC(Bielow et al., 2016) using the Maxquant result files.

The relative abundance of proteins was analysed by comparing their normalised label-free quantification (LFQ) intensities (Cox et al., 2014; Tyanova et al., 2016). Minimum LFQ values were calculated for each sample by taking the median of the 40 lowest LFQ values. Subsequently, LFQ values of non-detected proteins were replaced by the calculated minimum values. Afterwards, to correct for differences in total protein signal between samples, LFQ values were normalised by dividing each protein LFQ by the total protein LFQ of each sample. The normalised LFQ values were log_10_-transformed before unpaired two-tailed t-tests were performed, and fold changes were calculated. EggNOG-mapper v2 (Cantalapiedra et al., 2021) was used for functional annotation of proteins. The mass spectrometry proteomics data have been deposited to the ProteomeXchange Consortium via the PRIDE partner repository with the dataset identifier PXD074394 (Perez-Riverol et al., 2025)

### Electron microscopy

For electron microscopy analysis, *L. lactis* FM03P wild type and isolates were grown in a pH-controlled batch cultivation. Cells were harvested at the same timing as described in the proteomic analysis above (exponential and stationary phase) by centrifugation of 2 ml culture at 17000× g for 2 min at 4°C. Cell pellets were immediately resuspended in fixation buffer (2.5% glutaraldehyde in 0.1 M phosphate/citrate buffer). The samples were incubated for 1 h at room temperature and stored at 4 °C before further steps.

### Scanning electron microscopy (SEM)

The samples were washed 6 times in a washing buffer (0.1 M phosphate/citrate buffer). Then 1% osmium tetroxide in 0.1 M phosphate/citrate buffer was added and incubated for 1 h at room temperature, followed by 3 times washing in milliQ for 10 min. The samples were further dehydrated in a graded series of ethanol, followed by a critical point drying (CPD) with 100% ethanol. The dried samples were attached to aluminium stubs with double-sided carbon stickers and sputter coated with 12 nm tungsten. The samples were visualised using a scanning electron microscope (Magellan 400, FEI) at 2500 – 25000× magnification with secondary electron detection of 2.00 kV and 13 pA.

### Transmission electron microscopy (TEM)

The samples were washed 2 times with 0.1 M phosphate/citrate buffer, followed by resuspension in 100 µl of 3% gelatine in 0.1 M phosphate/citrate buffer and incubation for 20 min at 4°C until the gelatine was solidified. The samples were cut into pieces between 1 – 3 mm^3^ and fixed in 2.5% glutaraldehyde for 1 h, followed by 6 times washing for 10 min with 0.1 M phosphate/citrate buffer. Further, 1% osmium tetroxide was used for 1 h post-fixation, followed by 3 times washing in milliQ for 10 min. The samples were dehydrated in a graded series of ethanol and infiltrated with a gradually increasing concentration of resin in ethanol. After overnight incubation in 100% resin, the samples were polymerized at 70 °C for 8 h. Next, 50 nm section of samples were obtained using Leica EM UC7 microtome and further stained with uranyl acetate and lead citrate. The samples were visualized using JEOL JEM-1400 plus electron microscope (JEOL USA Inc., USA) at 120 kV.

### Growth at high osmotic pressure

At least 4 independent overnight cultures of *L. lactis* FM03P wild type and isolates were inoculated at 0.1% v/v into CDM (0.5% w/v lactose without nucleotides) supplemented with 0, 174, 348, 522, 696, and 870 mM of either potassium lactate (KLa) or potassium chloride (KCl), with or without 1 mM of glycine-betaine in 96-wells plates. The plates were incubated in a Versamax plate reader (Molecular Devices, Sunnyvale, CA, USA) at 30°C for 72 h. The OD was continuously measured every 10 min at 600 nm with 5 s shaking between reads. The measurement was automatically recorded in SoftMax Pro software (Molecular Devices). The maximum growth rate of the wild type and isolates in each condition was calculated with the R package “growthrates” (Petzoldt, 2020) using the OD data as input. The growth rate was determined at OD below 0.3.

### Amino acid analysis

The supernatant from proteomic samples were analysed for extracellular amino acids by ultrahigh-pressure liquid chromatography (UPLC) as described previously (Middendorf et al., 2022). Briefly, the samples were deproteinized with sulfosalicylic acid and derivatized by using the AccQ-Tag Ultra Derivatization kit (Waters, Milford, MA, USA). The extracellular arginine and ornithine concentration were determined by using Chromeleon software (Thermo Fisher Scientific).

### Structure model prediction and structural alignments

Structure models were predicted with AlphaFold2 (Jumper et al., 2021) using the RNAP β and β’ sequences of *Lactococcus* (WP_003130544.1 and WP_014570699.1) and sequences of mutants thereof (βV625 and β’R327). Inter- and intraprotein interactions of βV625 and β’R327 residues in the AlphaFold2 model were analysed using PDBe PISA v1.52 (Krissinel and Henrick, 2007). The closest structural homologs of the β/β’ dimer were identified using PDBeFold (Krissinel and Henrick, 2004). Structure models were aligned using the ‘super’ function in PyMOL (Schrödinger and DeLano, 2020). Figures of structure models were generated using PyMOL.

### Multiple sequence alignments

RNAP β and β’ sequences of *Lactococcus* (WP_003130544.1 and WP_014570699.1), *Bacillus subtilis* (6WVJ_2 and 6WVJ_3), *Mycobacterium tuberculosis* (8E8M_4 and 8E8M_5), *Mycobacterium smegmatis* strain ATCC 700084 (5VI8_3 and 5VI8_4), *Escherichia coli* strain K12 (6C9Y_2 and 6C9Y_3) and *Thermus aquaticus* (1I6V_2 and 2GHO_D) were aligned in Jalview (v2.11.3.2)(Waterhouse et al., 2009) using the Mafft alignment option (Nakamura et al., 2018) with default settings.

## Results

### *L. lactis* FM03P population dynamics during stressostat cultivation

To understand how *L. lactis* increased its lactate resistance during stressostat cultivations, metagenomes of two independent stressostat populations and genomes of isolates were sequenced at different time points. The two stressostat populations are illustrated with Muller plot (Fig. 1A). After only seven days of evolution, various genotypes were found in both stressostats, with mutations in DNA-directed RNA polymerase (RNAP) subunit β’ (*rpoC*) occurring in both stressostat cultivations. In addition, a mutation in *greA* (a RNAP-interacting transcription elongation factor) was found in stressostat 2. From 14 days onwards, mutations in DNA-directed RNA polymerase subunit β (*rpoB*) arose in both populations, as well as in the phosphate transport complex proteins *phoU* (stressostat 1) and *pstB* (stressostat 2). Additional genotypes accumulated at later time points (Figure 1C). These experiments reveal that the dominating RNAP complex mutations (*rpoC, rpoB,* and *greA*) never co-occur in the same isolate. Combined with the relative abundance of specific gene alleles (Fig. 1B), this indicates that at the end of the stressostat cultivations, the majority of both populations (∼70% of stressostat 1 and ∼90% of stressostat 2) had a mutation in one of the RNA polymerase subunits, while in stressostat 2, all isolates had an additional PstB:E206K mutation. Sequencing of individual isolates (Fig. 1C) showed that mutations in the phosphate transport complex (*phoU* and *pstB*) were found only in cells containing a *rpoB* mutation, which suggests that the *rpoB* mutation is required for the beneficial effect of mutations in *phoU* and *pstB*.

**Fig. 1.**
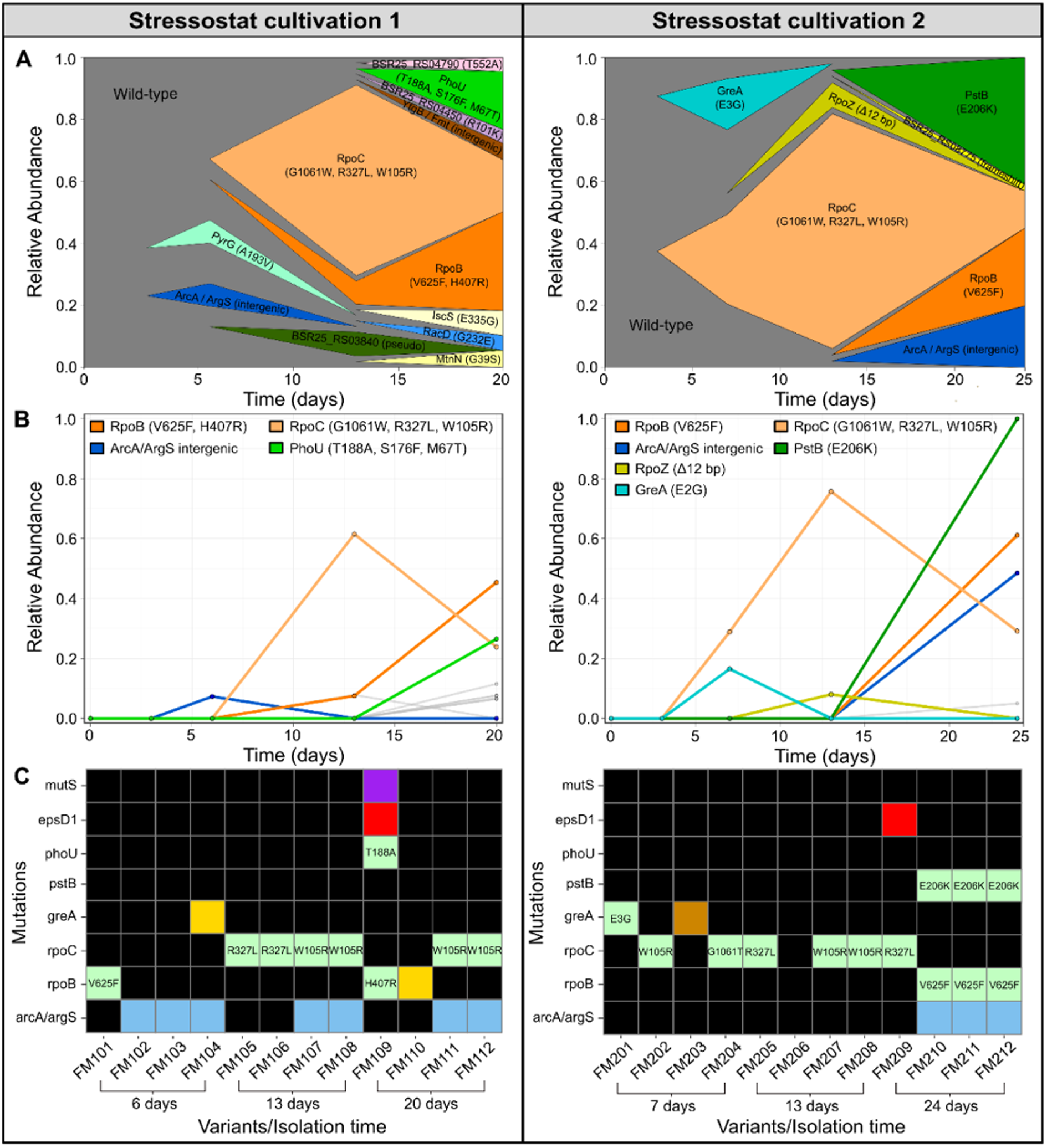
Dynamics of genomic mutations in L. lactis during two independent stressostat evolution experiments. (A) Muller plot representing the population dynamics in the stressostat cultivations determined by metagenomics. Colours indicate mutations in different genes. Labels indicate the exact mutations in the affected protein. (B) Relative abundance of specific genetic mutations determined by metagenomics. Only mutations that independently occurred in both stressostat cultivations are shown. (C) Mutations were found in the isolated mutants (Hartono et al., 2023) through whole genome sequencing and SNP analysis. Colours represent the effect of the mutation on the protein according to sequence ontology (SO) terms (green: missense variant; red: frameshift variant; yellow: conservative inframe insertion; brown: disruptive inframe deletion; purple: stop-gained, blue: non-coding region; black: no mutation). In Fig. 1B and 1C only mutations found in both stressostat cultivations are shown, and in addition mutS (hypermutator). The full list of gene alleles is provided in Table S1 and the full list of SNP analysis is provided in Figure S1 and Supplementary file 1.

Another mutation identified was located upstream of the gene encoding arginine deiminase (*arcA*). Aligning this sequence with other *L. lactis* strains revealed that the *arcA* genotype of the isolates is found in many other *L. lactis* wild-type strains. Based on this (meta)genomic analysis, we hypothesize that mutations in the RNAP complex improves fitness during the stressostat evolution experiments at high lactate concentrations. We selected *L. lactis* FM105 (further referred to as isolate RpoC:R327L) and *L. lactis* FM211 (further referred to as isolate RpoB:V625F) for further investigation as these genotypes were found in both stressostat populations. In addition, functional characterization of these isolates showed that RpoC:R327L produced more biomass than *L. lactis* FM03P (referred to as wild type (WT) isolate) when grown in a pH-controlled batch fermentation, and RpoB:V625F showed growth tolerance to higher lactate concentrations compared to the WT isolate (Hartono et al., 2023).

### Mutations in DNA-directed RNA polymerase subunit β (RpoB) and β’ (RpoC)

To obtain insights into the functional relevance of the mutations observed in the RNAP of *L. lactis* FM211 (RpoB:V625F) and FM105 (RpoC:R327L), we made multiple sequence alignments (MSAs) of the two subunits individually and structural alignments using a AlphaFold2-generated (Jumper et al., 2021) structural model of the *L. lactis* β/β’ heterodimer (Fig. 2).

**Fig. 2.**
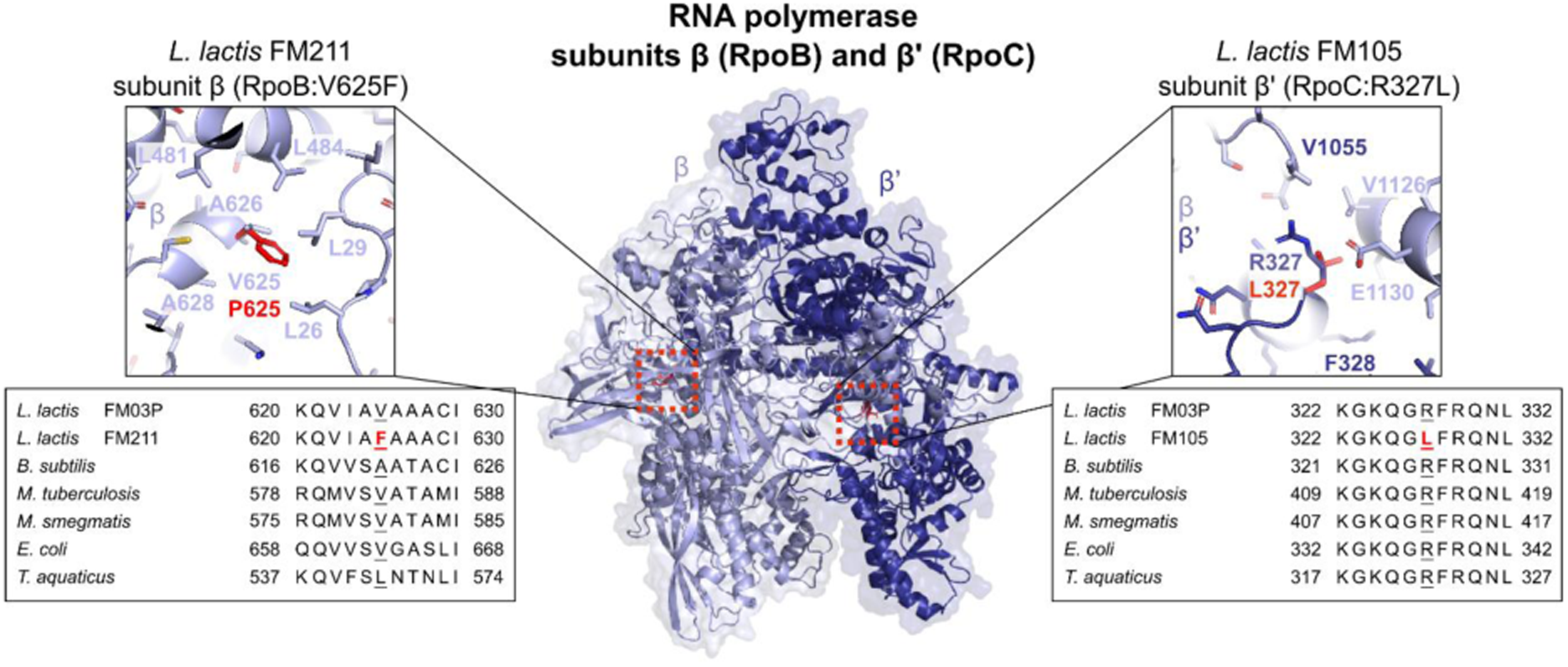
Implications of β (RpoB) V625F and β’ (RpoC) R327L substitutions. Middle: AlphaFold2-predicted model of the *L. lactis* FM03P RNAP subunits β (light blue) and β’ (dark blue) heterodimer. Please note that the RNAP complex contains additional components (Figure S2). Left: close-up view of β-subunit (RpoB) residue V625F substitution (red) in *L. lactis* FM211, and MSA of corresponding residues in RNAP homologs. Right: close-up view of β’-subunit (RpoC) residue R327L substitution (red) in *L. lactis* FM105, and MSA of corresponding residues in RNAP homologs.

RNAP comprises a α-subunit dimer, β and β’ subunits, an ω- subunit, and further requires sigma factors (σ) to initiate transcription(Cramer, 2002; Trinh et al., 2006; Lee et al., 2013). DNA-directed RNA polymerases (RNAP) complexes drive the transcription in all cellular organisms. Mutations in RNAP subunits can have important consequences. The *L. lactis* β/β’ dimer AlphaFold2 model is structurally highly similar to the β and β’ subunits of various bacterial holo-RNAP complexes (Table S2). The RpoB:V625F and RpoC:R327L mutations do not alter the predicted overall conformation of the β/β’ model (Fig. 2). In line with the mutant strains being viable, this suggests that these mutations are not disruptive for *L. lactis* RNAP structure and function.

At the position of WT *L. lactis* RpoB:V625, aliphatic residues are strictly conserved in homologous RNAP complexes (Fig. 2). RpoB:V625 is located in a hydrophobic pocket within the β subunit, which suggests that the hydrophobic aliphatic residues contribute to correct β subunit folding (Fig. 2). RpoB:V625 is not directly involved in interactions with other RNAP subunits, the DNA template, or the RNA product (Figure S2). However, RpoB:V625 is located only ∼10 Å away to a previously identified binding pocket for rifampicin, a RNAP inhibitor(Campbell et al., 2001). Although the mutant V625F residue is unlikely to directly interact with rifampicin, it might affect sensitivity to it. Indeed, growth experiments in presence of rifampicin show that the FM211 (RpoB:V625F) strain has an increased sensitivity to rifampicin (Figure S3).

At the position of *L. lactis* RpoC:R327, arginine residues are absolutely conserved in homologous RNAP complexes of all domains of life(Pupov et al., 2010) (Fig. 2). MSAs and comparison to other bacterial RNAP complex structures reveal that RpoC:R327 is located on the highly conserved Switch-2 (SW2) region within the β’ subunit (Fig. 2). SW2 is an important regulatory element that affects transcription initiation and RNAP processivity (Pupov et al., 2010). Consequently, it is likely that the RpoC:R327L mutation affects transcription. Modelling of the observed RpoB:V625F and ProC:R327L mutations suggests that these mutations affect RNAP-mediated transcription, conceivably resulting in proteome adjustments in these isolates.

### Proteomic responses in pH-controlled batch cultures

As the mutations in RpoC and RpoB could indirectly improve lactate resistance through resulting proteome changes, a comparative proteomic analysis was performed on WT and isolates during growth in pH-controlled batch fermentations. Samples were taken at the beginning of exponential phase (∼100 mM lactate in all three isolates) and at early stationary phase when biomass production stopped at ∼600 mM lactate for WT and RpoB:V625F, and at ∼700 mM lactate for RpoC:R327L (Figure S4).

To better understand general *L. lactis* stress response during pH-controlled batch fermentations, we first compared the proteome of the WT at the stationary phase versus the exponential phase (Fig. 3A). In the stationary phase, proteins related to the osmotic stress response, such as compatible solute uptake (*opp, kup, busAB*), amino acid synthesis (glutamate & proline), c-di-AMP regulon (*kup, busAB, opuAA, cbpE*), and cell wall synthesis proteins, were upregulated. However, despite higher abundance of these proteins, electron microscopy analysis of WT cells suggested cell damage from hyperosmotic condition indicated by cell lysis and protoplast formation at the stationary phase (Fig. 3B). The reduced efficacy of the osmotic stress defence might be a consequence of the absence of compatible solutes such as glycine-betaine (GB) in the defined medium used in the experiments. Additionally, proteins involved in the arginine deiminase pathway (ADI) (*arcA, arcC, argF*) were upregulated in the stationary phase. The ADI pathway converts arginine into ornithine, resulting in the generation of ATP and ammonia. Other upregulated proteins were related to oxidative stress, cation transport, the electron transport chain, and riboflavin synthesis, which suggests a redox imbalance exists.

**Fig. 3.**
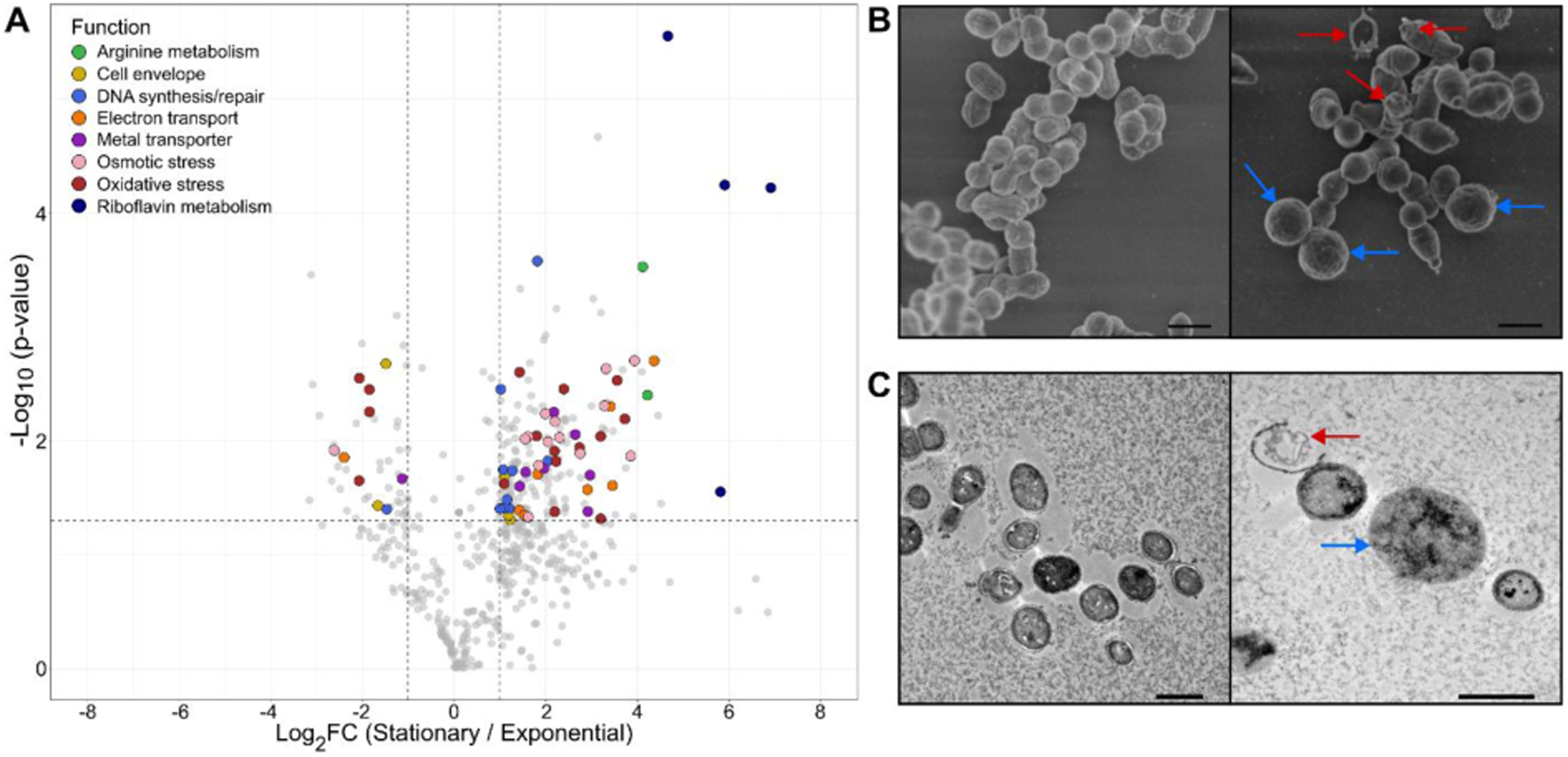
Proteomes and morphology changes of L. lactis FM03P during growth in pH-controlled batch fermentation. (A) Volcano plot of *L. lactis* FM03P proteomes of stationary growth phase (high lactate) vs exponential growth phase (low lactate). Positive log2-fold change indicates a higher protein abundance in the stationary phase than exponential phase, while negative value indicates a lower protein abundance. Dotted lines represent cut-offs for significance (at least 2-fold change and p-value ≤ 0.05, paired two-tailed t-test). Coloured circles indicated the proteins functions, while grey circles indicated proteins which were not investigated or of which abundance was not significantly altered. (B) Scanning electron microscopy (SEM) and (C) transmission electron microscopy (TEM) of *L. lactis* FM03P grown in pH-controlled batch cultures at the exponential phase (left) and stationary phase (right). Red arrows indicate damaged or lysed cells. Blue arrows indicate protoplasts. Scale bars represent 1µm.

Comparing the proteome of the WT with that of the isolates revealed similar adaptations in both isolates in exponential and stationary phase (Fig. 4 & Supp Fig 5). Expression of proteins involved in the ADI pathway were already upregulated in at the exponential phase in the isolates, compared to only at the stationary phase for the WT (Fig. 3A & Fig. 4). At the same time, no significant changes (below 2-fold threshold) were observed for majority of the other proteins (Fig. 4). This suggests that the ADI pathway was active in the isolates during the exponential phase, even before the carbon source was depleted. Notably, despite higher abundance of ADI pathway proteins in WT in the stationary phase (Fig. 3A), they were still more abundant in the isolates in this growth phase (Fig. 4). Higher abundance of ADI enzymes could be linked to increased arginine utilization following determination of extracellular concentrations of arginine and ornithine during pH-controlled batch culture (Fig. 5A). This was supported by the observation of elevated consumption of extracellular arginine at the exponential phase by both RpoB:V625F and RpoC:R327L and significantly higher extracellular ornithine pools as compared to what the WT isolate produced in the stationary phase (Fig. 5A). Notably, isolate RpoB:V625F fully consumed all extracellular arginine in the exponential phase, which might be related to an additional mutation in the upstream region of *arcA* in this isolate (Fig. 1C).

**Fig. 4.**
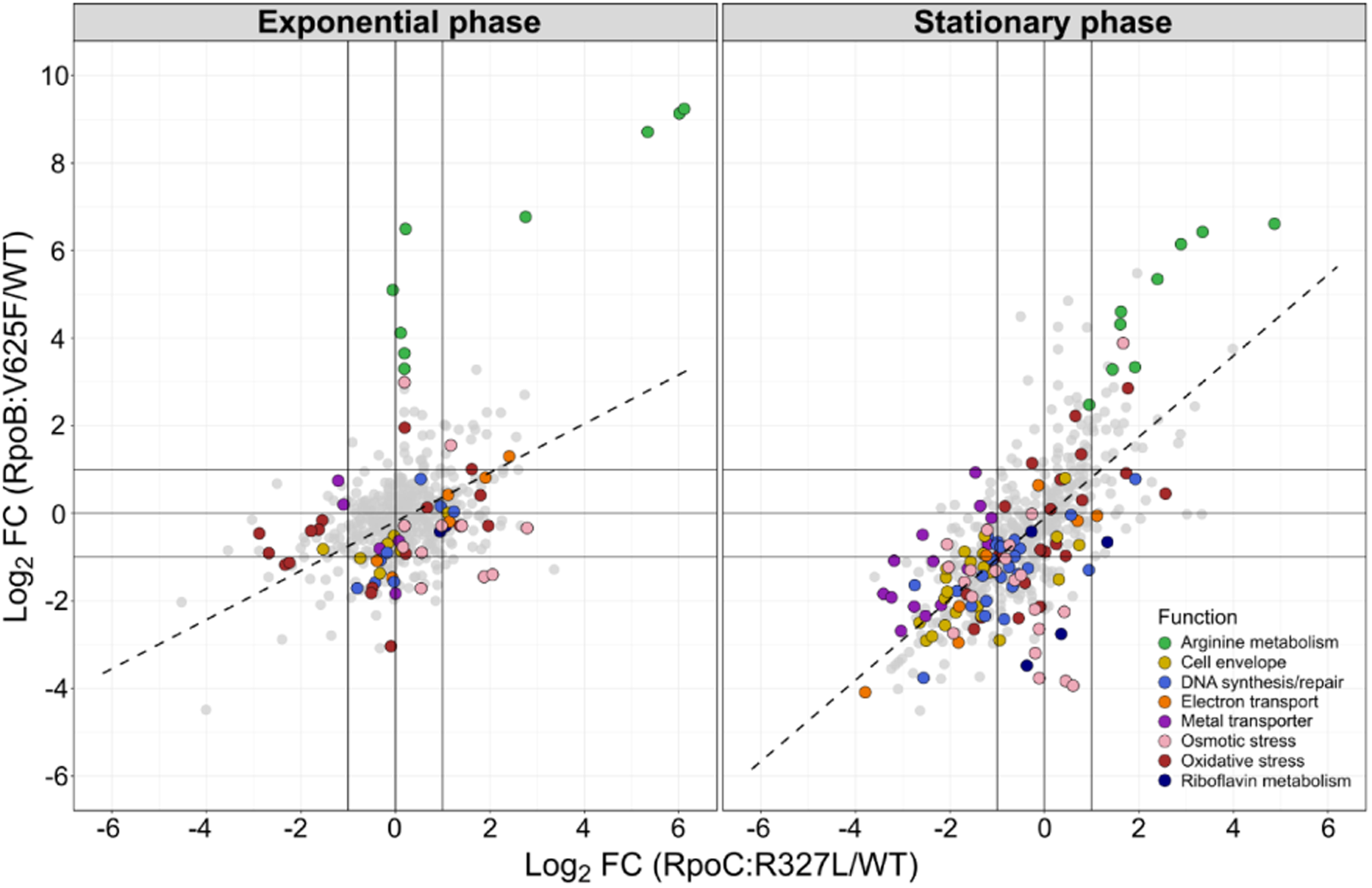
Proteome response of stressostat isolates compared to the WT at exponential (left) and stationary (right) phase. X-axis and Y-axis represent the log2-fold change of RpoC:R327 and RpoB:V625F compared to WT, respectively. Only significantly changed proteins in at least one isolate in either phase (unpaired two-tailed t-test, p-value ≤ 0.05) are shown in the plots. The coloured circles indicate relevant annotated protein functions, while grey circles indicate other proteins. Solid lines indicate a cut-off at 2-fold change.

**Fig. 5.**
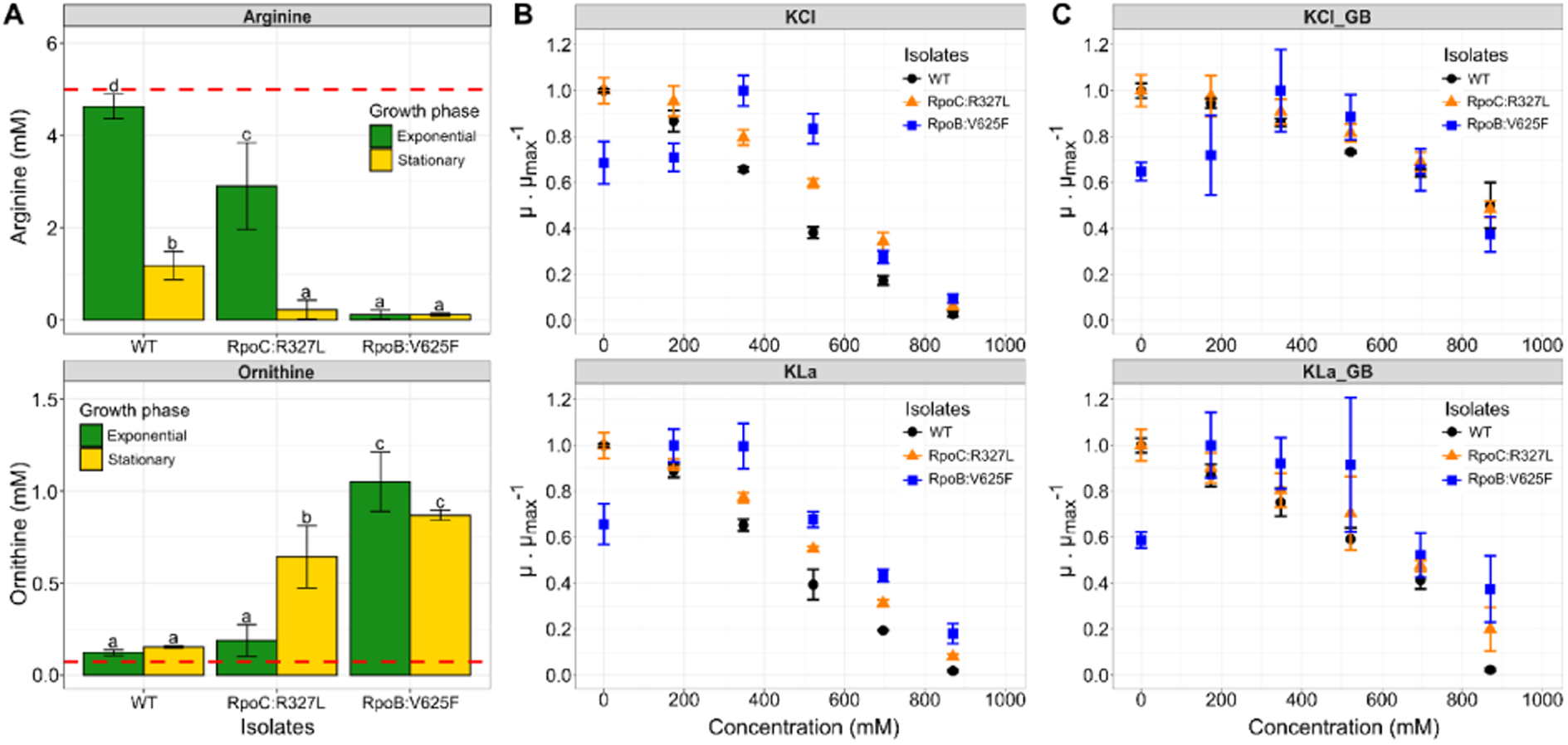
Phenotype of L. lactis FM03P and its stressostat isolates. (A) Extracellular arginine and ornithine concentrations during pH-controlled batch fermentations in exponential (green) and stationary (yellow) phase indicating the metabolism of arginine through the ADI pathway. Dotted red lines indicate the amino acid concentration in the blank CDM. Error bars represent the standard deviation of 3 biological replicates. Different letters indicate the significant difference in extracellular amino acid concentration (Dunn-Sidak posthoc, p ≤ 0.05). (B and C) Normalized growth rate at different potassium chloride (KCl; top) and potassium lactate (KLa; bottom) concentrations in batch cultures without pH-control in absence (A) or presence of glycine betaine (B). Colours represent the WT (black), RpoC:R327L (orange) and RpoB:V625F (blue). Error bars represent the standard deviation of 4 biological replicates. Statistical analysis is provided in Figure S7

In addition, other upregulated proteins in the WT isolate related to potential redox imbalance (oxidative stress, cation transport, electron transport chain and riboflavin synthesis), osmotic stress response, and cell wall synthesis were less abundant in the isolates compared to the WT in the stationary phase. In line with this observation, electron microscopy of the isolates revealed mostly intact cells in the stationary phase with similar morphology as exponential phase cells (Figure S6). These results indicate that the isolates suffered less from hyperosmotic stress despite being exposed to a similar lactate concentration as the WT (600-700 mM).

### Arginine metabolism and osmotic stress resistance

To confirm the enhanced hyperosmotic stress resistance of the isolates and to establish that high lactate concentration causes osmotic stress in *L. lactis*, the WT and isolates were grown in batch cultures in CDM with different concentrations of potassium chloride (KCl) or potassium lactate (KLa), in the absence or presence of glycine-betaine (GB), compatible solute known to alleviate osmotic stress (Graham and Wilkinson, 1992; Molenaar et al., 1993; van der Heide and Poolman, 2000; Verheul et al., 1997). The maximum growth rates were calculated from the growth curves for all conditions and presented as normalised growth rate against the solute concentration (Fig. 5B & Fig. 5C). This shows KLa inhibited growth similarly as KCl in the WT and isolates when no GB is added. Moreover, the isolates show higher growth rates than the WT at high salt (KCl and KLa) concentrations. This confirms that isolates were more resilient to high salt concentrations in the tested condition, which suggests that they are more resistant to osmotic stress and potential cell wall damage (Figure S6). Interestingly, the isolates showed higher resistance against high KLa concentrations compared to high KCl concentrations, whereas growth rates of the WT were the same for both salts. Supplementation with GB alleviated the osmotic stress caused by KCl and KLa in both WT and isolates (Fig. 5B). Growth rates of WT and isolates were not significantly different at high KCl concentration when supplemented with GB (Figure S7), but isolates RpoB:V625F did grow significantly faster than WT at the highest KLa concentration tested (850 mM). Interestingly, GB alleviated KCl inhibition much more than Kla inhibition, indicated by higher growth rates of WT and isolates in KCl than in KLa when GB was added. This suggests that although potassium lactate mainly causes osmotic stress, it might cause additional types of stress. Finally, isolate RpoB:V625F showed a possible evolutionary trade-off, with a significantly lower growth rate at low solute concentrations, indicating growth inhibition in hypoosmotic conditions. This isolate grew optimally when supplemented with ∼300 mM solutes. Combined, this shows that mutations in the RNA polymerase (RNAP) subunits in both isolates have various pleotropic effects on the proteome that result in increased osmotic stress resistance and ADI pathway activity.

## Discussion

In this study, we have elucidated underlying mechanisms of higher lactate resistance in *Lactococcus lactis* stressostat isolates with mutations in the RNA polymerase (RNAP) complex contributing to their dominance in evolved populations. Changes in transcription behaviour of RNAP can result in a pleotropic effect on the phenotypes, which can results in a benefit across multiple conditions or an evolutionary trade-off (Choudhury et al., 2023; Conrad et al., 2010; Jin and Gross, 1989; Trinh et al., 2006; Zhou and Jin, 1998). To the best of our knowledge, the RpoB:V625F mutation in the β subunit of RNAP has never been reported before. However, substitution of the hydrophobic valine with a bulky aromatic phenylalanine might affect RNAP interactions with DNA or other proteins and result in a stringent phenotype, which improves stress tolerance and long-term starvation in exchange of growth rate. Stringent mutations decrease stability in open complex intermediate initiation, leading to repression of growth-associated genes and activation of amino acids metabolism (Choudhury et al., 2023). The RpoB:V625F isolate indeed grew more slowly than WT *L. lactis* FM03P but had an increased arginine metabolism, which indicates that this RNAP mutation results in a trade-off with a stringent phenotype.

Meanwhile, the RpoC:R327L substitution is located in the highly conserved Switch-2 (SW2) region of the β’ subunit. The SW2 region is important for opening and closing of the RNAP cleft (Pupov et al., 2010). In homologous RNAP complexes, residues corresponding to R327 directly interact with template DNA and are important for DNA melting, but also for interactions with σ factors (Boyaci et al., n.d.; Dailler et al., 2021; Pupov et al., 2014, 2010). Therefore, mutation in RNAP at R327 is expected to change RNAP transcription behaviour. It has previously been reported that an increased stringent phenotype can be achieved without having a trade-off in growth phenotype when the mutations in RNAP result in intermolecular polar interactions and decreased target-Bridge Helix interactions (Choudhury et al., 2023). While the exact molecular mechanism underlying the effects of the RpoC:R327L substitution remains unclear, we observed no growth trade-off in this isolate despite the increased lactate resistance and higher arginine metabolism.

The wide range of effects from β and β’ mutations make them a hotspot for adaptive mutations in RNAP. The mutations affecting global transcription patterns appears to be common outcome of ALE experiments in fluctuating stress conditions, as they provide greater flexibility for survival and improved fitness during evolutionary experiments (Choudhury et al., 2023; Trinh et al., 2006). RNAP mutants isolated from our ALE experiment with particular conditions and stresses are known to also provide protection against other stresses (Conrad et al., 2010; Dragosits et al., 2013). Whereas mutations in RpoC were already found after three days of incubation in the stressostat, mutations in RpoB only appeared later in the ALE at higher lactate concentrations. Interestingly, RpoC and RpoB mutations were never found simultaneously, as shown in the WGS of the isolates. Instead, RpoB mutants replaced the RpoC mutants in the stressostat populations. As RNAP regulates the global DNA transcription in bacteria, mutations in both subunits might affect the integrity of the RNAP and could be detrimental to the fitness of the isolate. It appears that RNAP mutations provide initial adaptation with a broad effect, while additional mutations further enhance fitness in that specific condition (Applebee et al., 2008; Herring et al., 2006). However, a compensatory mutation could occur within the same or in a different RNAP subunit (Comas et al., 2011; Brandis et al., 2012; de Vos et al., 2013). A RpoB rifampicin-resistant mutant was evolved through serial propagation in rich media with rifampicin, selecting for a higher growth rate. This mutant had mutations in either RpoA, RpoB or RpoC (Brandis et al., 2012). These double mutants maintain resistance to rifampicin but had a higher growth rate. It seems that the compensatory mutations restored the properties of RNAP that were altered by the initial mutation (Brandis et al., 2012). Finally, in our study, we found different mutations in selected isolates within the same subunit. Notably, in the tested condition, all isolates exhibited higher lactate resistance than wild type *L. lactis* FM03P (Hartono et al., 2023), indicating that different RNAP mutations allow *L. lactis* FM03P to cope with increasing lactate stress in the stressostat cultivation.

In the stationary phase, *L. lactis* FM03P wild type (WT) upregulated mainly proteins related to osmotic stress resistance as a response to lactate stress in a pH-controlled batch culture in the absence of compatible solute GB. Ntably, RNAP mutations changed isolates’ responses to lactate stress in this condition. The isolates’ proteome exhibited a significantly lower abundance of proteins associated with the osmotic stress response compared to the WT, which included compatible solute transporters (glycine-betaine), potassium transporters and proteins involved in proline synthesis. Expression of potassium (*kup*) and compatible solute (*opuAA, busAB*) transporters is regulated by c-di-AMP, which is a global osmo-regulator in many Gram-positive bacteria (Pham et al., 2021; Turner et al., 2023). However, as glycine-betaine was not supplemented in the growth media, this upregulation of the glycine-betaine transporters in the WT cannot be considered as an effective stress response. Additionally, both isolates conceivably maintain a lower c-di-AMP pool than WT due to lower abundance of c-di-AMP synthesis proteins and receptor proteins (Supplementary file 2). Lack or absence of c-di-AMP can cause toxicity on bacterial cell (Choi et al., 2015; Devaux et al., 2018; Pham et al., 2021). All RpoB isolates acquired mutations in the phosphate transport complex (*phoU, pstB*) at the end of stressostat cultivations (Fig. 1C). The phosphate transport complex is important for cell survival when c-di-AMP is downregulated. Compensatory mutation in *pstB* was reported to attenuate the toxicity of *busAB* (glycine-betaine transporter) when c-di-AMP is lacking (Devaux et al., 2018). Furthermore, proteins involved in proline synthesis from peptides and glutamate (GltA, ProC) were more abundant in the WT compared to the isolates in the stationary phase. Proline is a compatible solute that is taken up and synthesised in high salinity environments (Molenaar et al., 1993; Brill et al., 2011; Hoffmann et al., 2012). We detected a significantly higher extracellular proline concentration for the WT than the isolates (Figure S8). We hypothesise that a higher extracellular proline concentration may indicate a higher intracellular proline concentration in the WT than in the isolates. This could be linked to higher proline synthesis and/or release from peptides in the WT, followed by leakage of proline from damaged and/or lysed cells.

Despite all these osmotic stress responses, growth of the WT was still severely inhibited at high lactate concentrations without GB. In contrast, the isolates exhibited higher osmotic stress resistance than the WT, but lower expression of these proteins, which are commonly known as resistance mechanisms against osmotic stress. Instead, the isolates contained lower levels of some proteins related to peptidoglycan synthesis such as MurE involved in the cross-linking of peptidoglycan (Garde et al., 2021). Downregulation of MurE may reduce or modify the cross-linking structure in the peptidoglycan which can increase osmotic stress resistance (Piuri et al., 2005). Although we did not observe a noticeable change in cell wall structure in TEM pictures (Figure S6), both isolates showed a higher sensitivity to cefuroxime, an antibiotic targeting the bacterial cell wall (Figure S9), indicating a possible more open, less rigid peptidoglycan structure compared to the WT. A more detailed analysis of cell wall modifications in the isolates can confirm whether such adjustments are an effective mechanism to cope with osmotic stress in pH-controlled batch cultivation, especially when external compatible solutes such as GB are not present.

Finally, we showed that lactate stress mainly causes osmotic stress, indicated by the proteome responses and the similar growth inhibition as KCl, in line with previous research (Loubiere et al., 1997). Moreover, during the pH-controlled batch cultivation, potassium hydroxide (KOH) was added to maintain the desired pH level. The resulting K^+^-ions contribute to the increase in osmotic pressure during the fermentation, which theoretically increases approximately 8-fold (1 lactose α 4 lactate^-^ + 4 K^+^). However, GB did not alleviate lactate inhibition to the same extent as KCl inhibition (Fig. 5C). In addition, there was a discrepancy in the growth rate of *L. lactis* in pH-controlled batch cultures compared to batch cultures without pH control with supplemented sodium lactate (Loubiere et al., 1997). This suggests that lactate accumulation causes more than osmotic stress, especially in pH-controlled batch cultivations. In the presented proteomes, proteins involved in oxidative stress response, electron transport chain, cation transporters, and riboflavin synthesis were upregulated in the stationary phase of the WT and significantly less abundant in the isolates. Therefore, we hypothesize that lactate accumulation in absence of GB may also cause a redox imbalance, which causes the upregulation of the aforementioned proteins. Further research is needed to test this hypothesis.

### Conclusion

In conclusion, this study revealed that *L. lactis* improved its lactate resistance during stressostat cultivation initially by mutations in the RNA polymerase β and β’ subunits, altering the proteome. This increased the hyperosmotic stress resistance of the isolates in absence of GB, possibly by modulation of crosslinking of peptidoglycan in the cell wall. Subsequently, other mutations appeared in the stressostat population, such as a SNP in the upstream region of *arcA* activating the ADI pathway to generate additional ATP, and mutations in the phosphate transport complex (*pstB*), conceivably to attenuate the toxicity of c-di-AMP downregulation. While lactate accumulation primarily causes osmotic stress to the bacteria, similar to salt, proteome changes suggest that lactate may also (indirectly) induce oxidative stress. This highlights that RNAP mutations cause pleiotropic effects to alleviate complex stresses caused by high lactate concentrations.

## CRediT authorship contribution statement

SH: Conceptualisation, Data curation, Formal analysis, Investigation, Methodology, Visualisation, Writing-original draft and editing; HLR: Formal analysis, Writing - Review & Editing; SB: Methodology, Investigation, Formal analysis; DCS: Visualisation, Formal analysis, Writing - Review & Editing; TA: Conceptualisation, Funding acquisition, Supervision, Writing - Review & Editing; EJS: Conceptualisation, Project administration, Funding acquisition, Supervision, Writing - Review & Editing; OVM: Conceptualisation, Project administration, Funding acquisition, Supervision, Formal analysis, Visualisation, Writing - Review & Editing.

## Acknowledgement

We thank Jelmer Vroom (Wageningen Electron Microscopy Centre, Wageningen University & Research) for his assistance in SEM and TEM.

## Data availability

Metagenomic sequencing datasets can be accessed at ENA: PRJEB108644. Whole genome sequencing datasets can be accessed at ENA: PRJEB108642. The mass spectrometry proteomics datasets can be accessed at PRIDE: PXD074394

