## Supplementary materials for "Stressostat cultivation of *Lactococcus lactis* improves lactate stress resistance through mutations in RNA polymerase"

**Table S1. Allele frequency of L. lactis FM03P communities in two independent stressostat cultivations.**

| **Description** | **Mutation** | **Annotation** | **Gene** | **Position** | **Stressostat cultivation 1 (days)** | | | | | **Stressostat cultivation 2 (days)** | | | | |
| --- | --- | --- | --- | --- | --- | --- | --- | --- | --- | --- | --- | --- | --- | --- |
|  |  |  |  |  | **0** | **3** | **6** | **13** | **20** | **0** | **3** | **7** | **13** | **24** |
| 5'‑methylthioadenosine/adenosylhomocysteine nucleosidase | C→T | G39S (GGC→AGC) | *mtnN* ← | 1453753 |  |  |  |  | 7.70% |  |  |  |  |  |
| acetolactate synthase large subunit | C→T | pseudogene (1009/1728 nt) | *BSR25_RS03840* → | 754287 |  |  |  | 7.80% |  |  |  |  |  |  |
| arginine deiminase/arginine‑‑tRNA ligase | G→A | intergenic (‑56/+246) | *arcA* ← / ← *argS* | 1612627 |  |  | 7.30% |  |  |  |  |  |  | 48.50% |
| aspartate/glutamate racemase family protein | C→T | G232E (GGA→GAA) | *racD* ← | 1839089 |  |  |  |  | 6.80% |  |  |  |  |  |
| CTP synthase | C→T | A193V (GCC→GTC) | *pyrG* → | 29647 |  |  | 7.40% |  |  |  |  |  |  |  |
| cysteine desulfurase | A→G | E335G (GAA→GGA) | *iscS* → | 1432553 |  |  |  |  | 11.50% |  |  |  |  |  |
| DNA‑directed RNA polymerase subunit beta | C→A | V625F (GTT→TTT) | *rpoB* ← | 1366405 |  |  |  | 7.50% | 19.80% |  |  |  |  | 61.10% |
| DNA‑directed RNA polymerase subunit beta | T→C | H407R (CAC→CGC) | *rpoB* ← | 1367058 |  |  |  |  | 25.60% |  |  |  |  |  |
| DNA‑directed RNA polymerase subunit beta' | C→A | G1061W (GGG→TGG) | *rpoC* ← | 1361398 |  |  |  | 15.20% | 5.60% |  |  | 6.90% | 9.70% |  |
| DNA‑directed RNA polymerase subunit beta' | C→A | R327L (CGT→CTT) | *rpoC* ← | 1363599 |  |  |  | 15.60% |  |  |  | 13.90% | 49.40% | 29.10% |
| DNA‑directed RNA polymerase subunit beta' | A→G | W105R (TGG→CGG) | *rpoC* ← | 1364266 |  |  |  | 30.70% | 18.20% |  |  |  |  |  |
| DNA‑directed RNA polymerase subunit beta' | A→T | W105R (TGG→AGG) | *rpoC* ← | 1364266 |  |  |  |  |  |  |  | 8.10% | 16.60% |  |
| DNA‑directed RNA polymerase subunit omega | Δ12 bp | coding (287‑298/354 nt) | *rpoZ* ← | 1469693 |  |  |  |  |  |  |  |  | 8.00% |  |
| GlsB/YeaQ/YmgE family stress response membrane protein/methionyl‑tRNA formyltransferase | A→G | intergenic (‑69/+106) | *ytgB* ← / ← *fmt* | 1464700 |  |  |  |  | 7.60% |  |  |  |  |  |
| NAD(P)/FAD‑dependent oxidoreductase | +A | coding (287/1332 nt) | *BSR25_RS08775* ← | 1725696:1 |  |  |  |  |  |  |  |  |  | 5.00% |
| NusG domain II‑containing protein | G→A | R101K (AGG→AAG) | *BSR25_RS04450* → | 886009 |  |  |  |  | 6.40% |  |  |  |  |  |
| phosphate ABC transporter ATP‑binding protein | C→T | E206K (GAA→AAA) | *pstB* ← | 1274944 |  |  |  |  |  |  |  |  |  | 100% |
| phosphate signaling complex protein PhoU | T→C | T188A (ACA→GCA) | *phoU* ← | 1273099 |  |  |  |  | 11.50% |  |  |  |  |  |
| phosphate signaling complex protein PhoU | G→A | S176F (TCT→TTT) | *phoU* ← | 1273134 |  |  |  |  | 8.90% |  |  |  |  |  |
| phosphate signaling complex protein PhoU | A→G | M67T (ATG→ACG) | *phoU* ← | 1273461 |  |  |  |  | 6.10% |  |  |  |  |  |
| PTS glucose transporter subunit IIA | T→C | T552A (ACT→GCT) | *BSR25_RS04790* ← | 956273 |  |  |  |  | 6.50% |  |  |  |  |  |
| transcription elongation factor GreA | A→G | E3G (GAA→GGA) | *greA* → | 166802 |  |  |  |  |  |  |  | 16.50% |  |  |

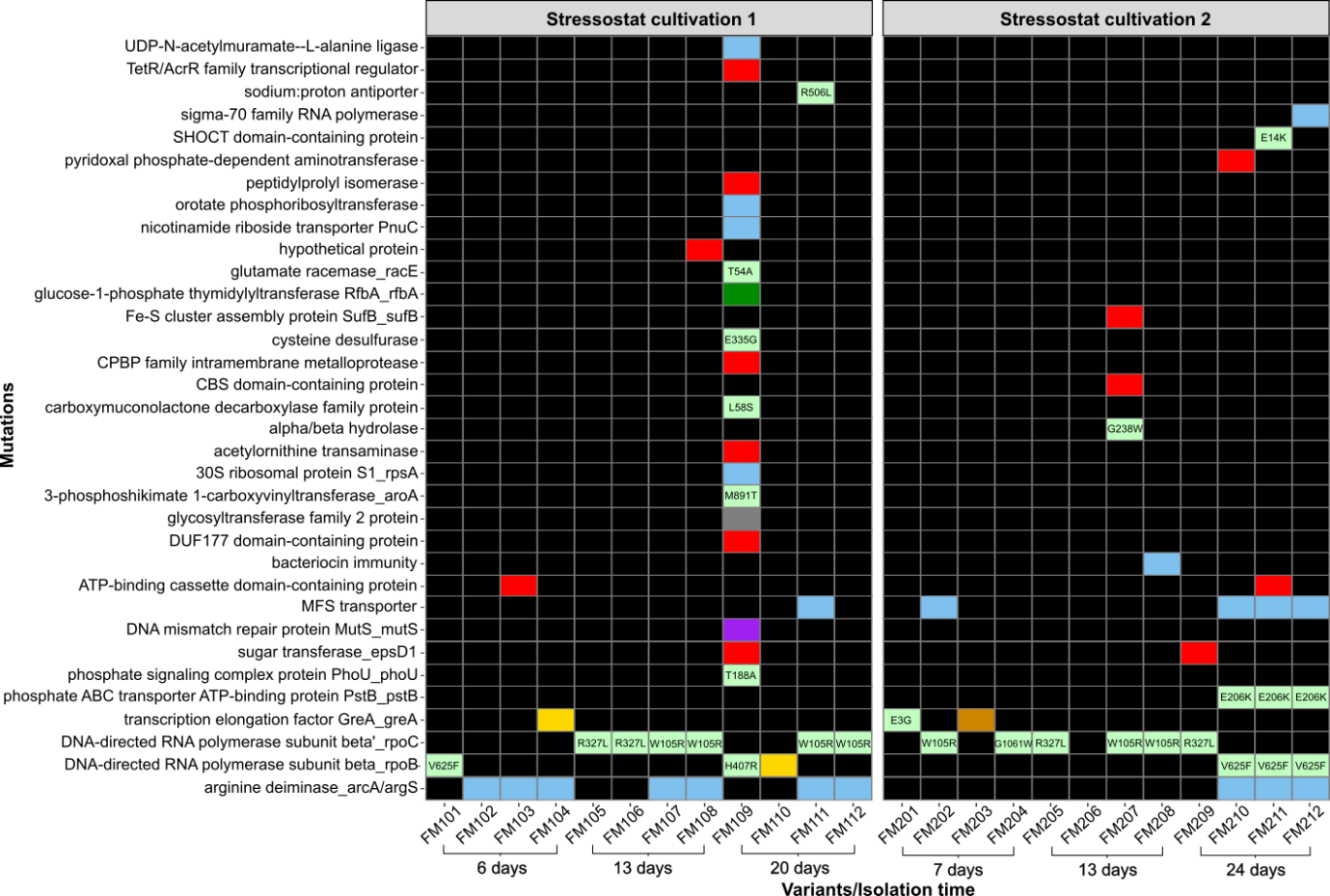

**Figure S1. All mutations found in the isolated variants through SNP analysis.** The variants were isolated from two independent stressostat evolution. Colours represent the effect of the mutation on the protein according to sequence ontology (SO) terms (light green: missense variant; red: frameshift variant; yellow: conservative in-frame insertion; brown: disruptive in-frame deletion; purple: stop-gained, green: synonymous variant; grey: frameshift variant & start lost; blue: non-coding region; black: no mutation)

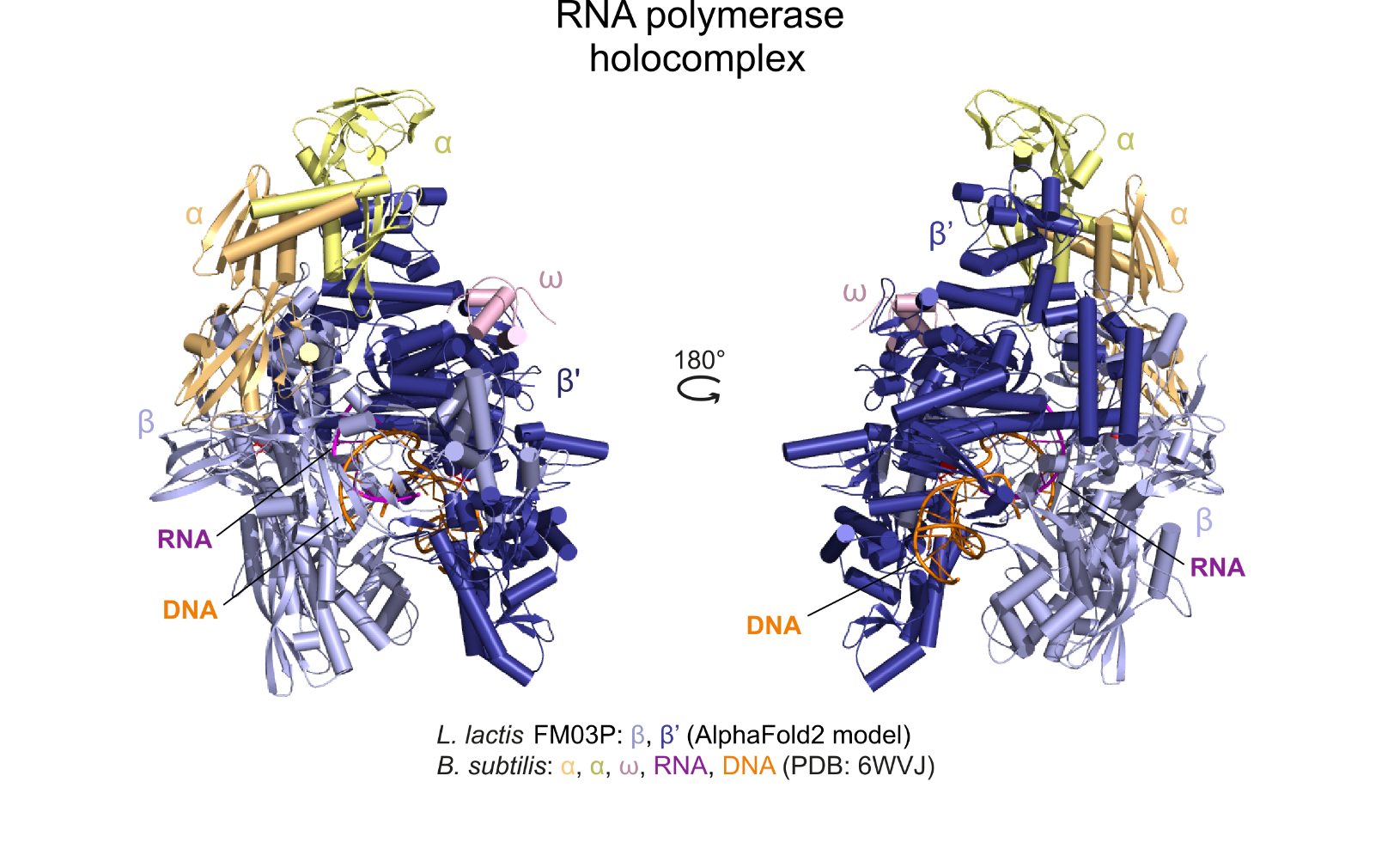

**Figure S2. General composition of the RNAP holocomplex.** The AlphaFold2-predicted model of the Lactococcus lactis FM03P RNAP subunits β (light blue) and β’ (dark blue) heterodimer was superposed on the RNAP structure of Bacillus subtilis (PDB: 6WVJ; RMSD: 1.0 Å over 13074 atoms). In the figure, β (light blue) and β’ (dark blue) subunits of L. lactis strain FM03P, and α (light orange and yellow) and ω (pink) subunits, as well as bound RNA (magenta) and DNA (dark orange) ligands of the aligned B. subtilis RNAP complex are displayed.

**Table S2. Structural similarity of L. lactis FM03P β/β’ dimer to the β and β’ subunits of various bacterial holo-RNAP complexes**

| **Species** | **PDB** | **RSMD** |
| --- | --- | --- |
| *Bacillus subtilis* | 6WVJ | 1.0 Å over 13074 atoms |
| *Mycobacterium smegmatis* | 5VI8 | 1.3 Å over 12072 atoms |
| *Mycobacterium tuberculosis* | 8E8M | 1.3 Å over 11941 atoms |
| *Eschericia coli* | 5UAH | 1.8 Å over 11717 atoms |
| *Thermus aquaticus* | 1I6V | 2.1 Å over 9373 atoms |

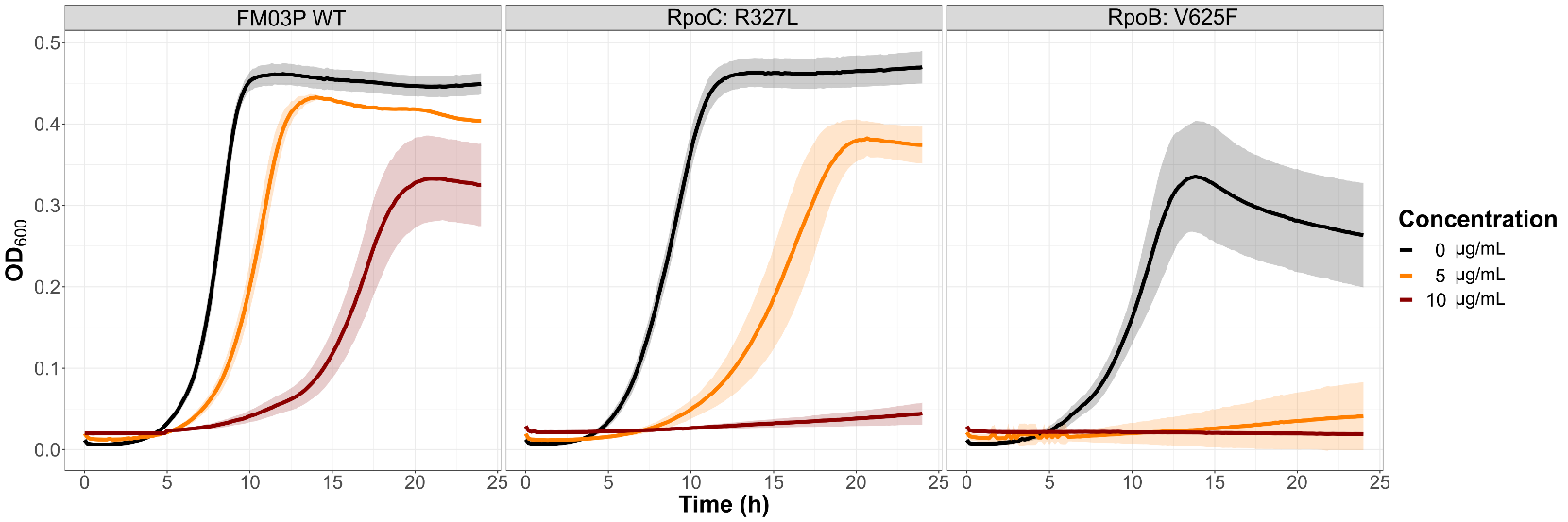

**Figure S3. Growth of L. lactis FM03P WT, RpoC:R327L, and RpoB:V625F in CDM with rifampicin**. Colours indicate rifampicin concentration: 0 (black), 5 (orange), and 10 (red) µg/mL rifampicin. Solid lines indicate the average and shadings indicate the standard deviation of OD_600_ of 4 biological replicates.

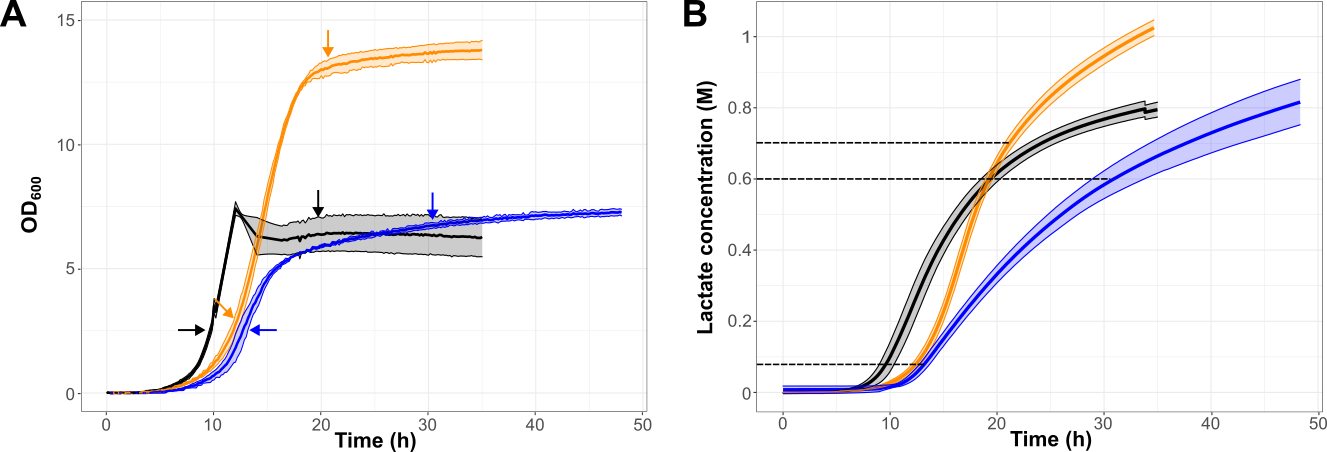

**Figure S4. Growth of L. lactis FM03P WT (black), RpoC:R327L (orange), and RpoB:V625F (blue) in pH-controlled batch cultures.** **A**: Biomass measured by OD_600_. **B**: lactate formation calculated from a linear correlation with KOH addition. Solid lines represent the averages and shadings represent the standard deviations of 3 biological replicates. Arrows (A) and dotted lines (B) indicate the timing of sampling for proteomic analysis.

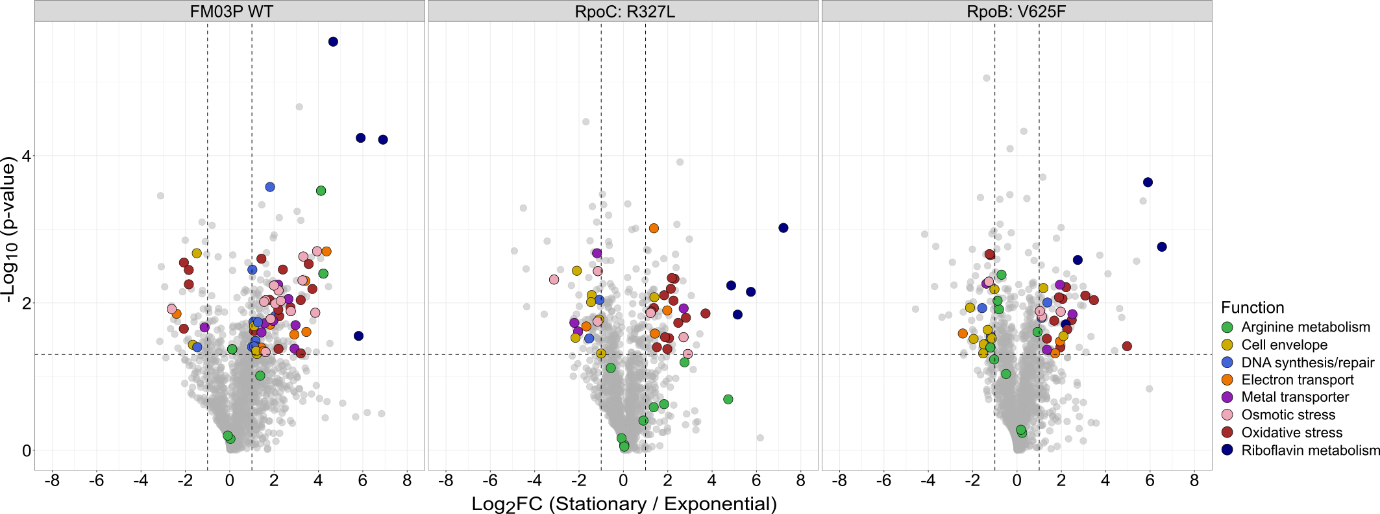

**Figure S5. Proteome changes of L. lactis FM03P and isolates during growth in pH-controlled batch fermentation.** Volcano plot of L. lactis FM03P WT (left), RpoC:R327L (middle), and RpoB:V625F (right) of stationary phase (high lactate) vs exponential phase (low lactate). Positive log2 fold change indicates a higher protein abundance in the stationary phase than exponential phase, while negative value indicates a lower protein abundance. Dotted lines represent cut-offs for significance (at least 2-fold change and paired two-tailed t-test, p-value ≤ 0.05, paired two-tailed t-test). Coloured circles indicated the proteins functions, while grey circles indicated uncharacterized and not significantly changed proteins.

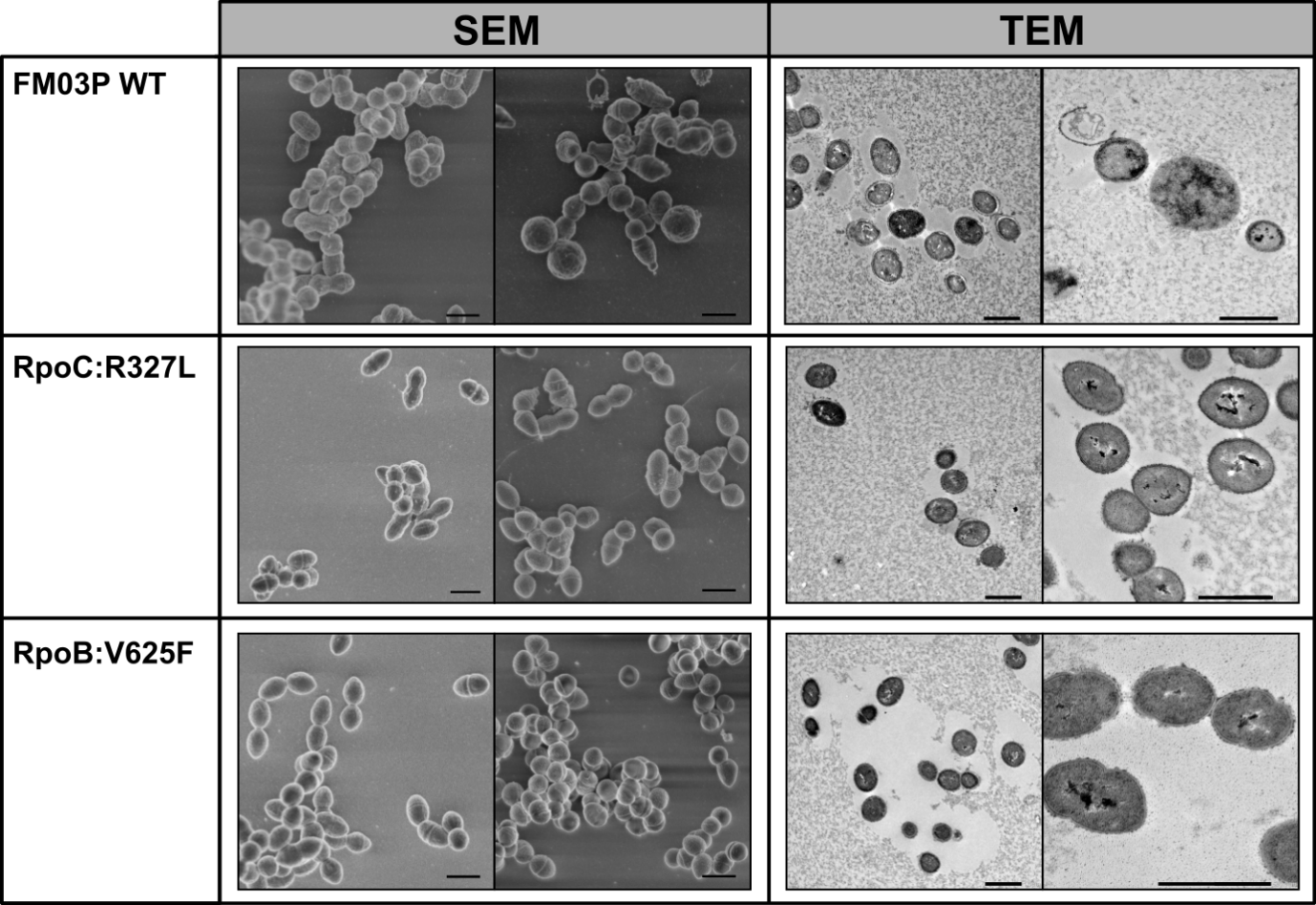

**Figure S6. Scanning electron microscopy (SEM) and transmission electron microscopy (TEM) of L. lactis FM03P WT, RpoC:R327L, and RpoB:V625F during growth in pH-controlled batch fermentation.** Cell morphology at different growth phase, exponential (left) and stationary (right) are shown in the figure for each method. Scale bars represent 1 µm.

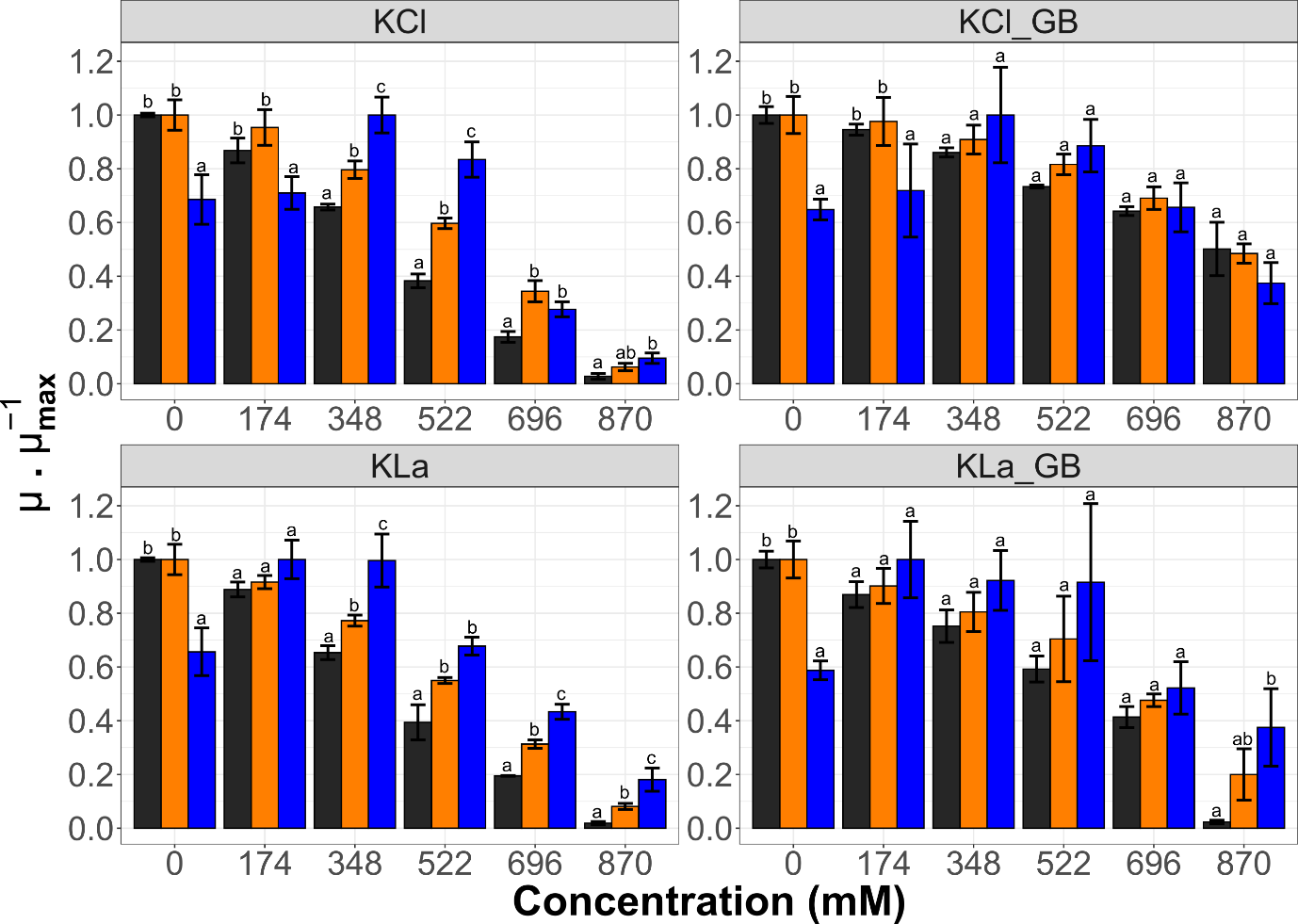

**Figure S7. Normalized growth rate at different potassium chloride salt (KCl; top) and potassium lactate (KLa; bottom) concentrations in batch cultures without pH-control in absence (left) or presence of glycine betaine (right).** Colours represent the WT (black), Rpoc:R327L (orange) and RpoB:V625F. Error bars represent the standard deviation of 4 biological replicates. Different letters indicate a significant difference in normalized growth rate between variants and WT within the same solute concentration (Dunn-Sidak posthoc, p ≤0.05).

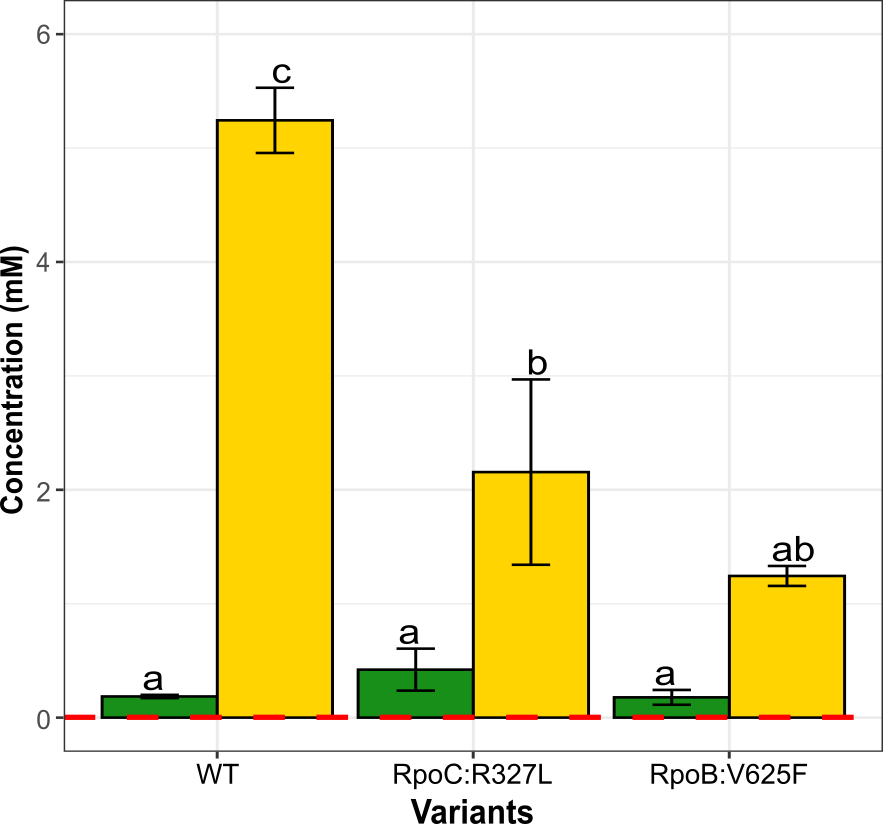

**Figure S8. Extracellular proline concentration of L. lactis FM03P WT, RpoC:R327L, and RpoB:V625F** **in pH-controlled batch cultures.** Colours indicate different growth phases: exponential (green) and stationary (yellow). The dotted red line indicates the proline concentration in the blank CDM. Error bars represent the standard deviation of 3 biological replicates. Different letters indicate a significant difference in extracellular proline concentration (Dunn-Sidak posthoc, p ≤0.05).

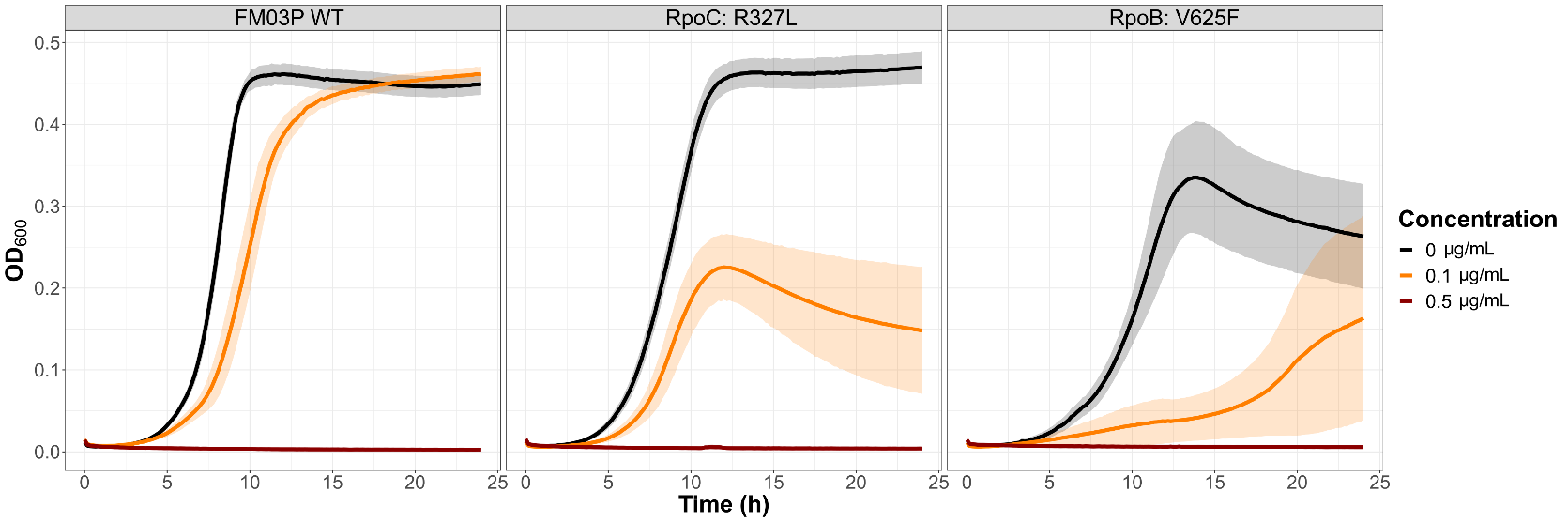

**Figure S9. Growth of L. lactis FM03P WT, RpoC:R327L, and RpoB:V625F in CDM with cefuroxime.** Colors indicate cefuroxime concentration: 0 (black), 0.1 (orange), and 0.5 (red) µg/mL of cefuroxime. Solid lines indicate the average, and shadings indicate the standard deviation of OD_600_ of 4 biological replicates.
